# Humanization of wildlife gut microbiota in urban environments

**DOI:** 10.1101/2022.01.05.475040

**Authors:** Brian A. Dillard, Albert K. Chung, Alex R. Gunderson, Shane C. Campbell-Staton, Andrew H. Moeller

**Affiliations:** Department of Ecology and Evolutionary Biology, Cornell University, Ithaca, NY 15853, USA; Tulane University, New Orleans, LA 70118, USA; Princeton University, Princeton, NJ 08544, USA

## Abstract

Urbanization is rapidly altering Earth’s environments, demanding investigations of the impacts on resident wildlife. Here, we show that urban populations of coyotes (*Canis latrans*) and crested anole lizards (*Anolis cristatellus*) acquire gut microbiota constituents found in humans, including the gut bacterial lineages most significantly associated with urbanization in humans (e.g., *Bacteroides*). Comparisons of urban and rural wildlife and human populations revealed significant convergence of the gut microbiota among urban host populations. Remarkably, all microbial lineages found in humans that were overrepresented in urban wildlife relative to rural wildlife were also overrepresented in urban humans relative to rural humans. These results indicate parallel effects of urbanization on human and wildlife gut microbiota and suggest spillover of bacteria from humans into wildlife in cities.

---

The gut microbial communities of vertebrates tend to reflect their host’s phylogenetic histories. Across a diversity of vertebrate clades, the community composition of the gut microbiota is on average more similar within host species than between host species, and microbiota dissimilarity between host species is positively associated with host evolutionary divergence time (*1*–*6*). However, anthropogenic influences can disrupt the host-lineage specificity of vertebrate gut microbiota through a process of humanization, in which hosts acquire gut microbiota constituents found in humans (*7, 8*), with potentially adverse consequences for host phenotypes and fitness. For example, captive mammals can acquire human gut microbiota constituents, creating mismatches between host and microbiota evolutionary history that have been implicated in the gastrointestinal disorders often experienced by captive hosts (*7*, *8*). Experiments in gnotobiotic rodents have further indicated that seeding hosts with human gut microbiota (or other non-native gut microbiota from a distantly related host species) can lead to stunted immunological development and reduced growth rates relative to seeding with native gut microbiota (*9*, *10*). With the influence of humans on ecosystems becoming more pronounced globally, there is a need to better understand the effects of the Anthropocene on vertebrate wildlife, including the possibility of humanization of wildlife microbiota. Of particular importance are the effects of urbanization, which are escalating at accelerating rates (*11*, *12*). Urbanization can alter the composition of the gut microbiota in diverse species of vertebrate wildlife (*13*, *14*), leading to consistent differences between populations in urban and rural settings. However, whether urbanization promotes the humanization of the microbiota of resident wildlife has not yet been explored.

To address this issue, we compared the gut microbiota of two vertebrate species—crested anoles (*Anolis cristatellus*; hereafter “anoles”) and coyotes (*Canis latrans*)—in both urban and rural settings to the gut microbiota of humans living in urban and rural settings. Anoles and coyotes maintain both urban and rural populations throughout North America and have become model systems for understanding the impacts of urbanization on vertebrate biology (*15*, *16*). We analyzed 492 fecal microbiota profiles from 94 anoles, 78 coyotes, and 320 adult humans (Supplemental Materials). Sampling locations for anoles included the city of Mayagüez on the western coast of Puerto Rico and an eastern longitudinal transect through Quemado into the rural areas of Maricao (Figure 1—figure supplement 1). Locations for coyotes included the city of Edmonton, Canada and the more rural areas around the neighboring small city of Leduc (Figure 1—figure supplement 1) (*14*). Locations for humans included urban centers across the United States, rural communities in Malawi, and rural Amazonian villages in Venezuela (*17*). Metadata for all samples are presented in Table S1. Across these samples we observed 29,492 unique Amplicon Sequence Variants (ASVs). The relative abundances of ASVs are presented in Table S2. Taxonomic assignments for all ASVs against the Silva 138 database are presented in Table S3. Alpha diversities and taxonomic barplots for all sample groups are presented in Figure 1—figure supplement 2 and Figure 1—figure supplement 3, respectively.

To assess the hypothesis that gut microbiota of wildlife living in urban settings have converged with those of humans relative to wildlife living in more rural settings, we calculated all microbiota dissimilarities (Binary Sorensen-Dice and Bray-Curtis) between pairs of samples using QIIME2 (*18*). PERMANOVA and PERMDISP indicated significant differences in gut microbiota composition among wildlife populations in different sampling locations (Supplemental Materials). To visualize the degree of compositional similarity among the gut microbiota profiles from anoles, coyotes, and humans, we next generated non-metric multidimensional scaling (NMDS) plots of Bray-Curtis and binary Sorensen-Dice dissimilarities. Results of these analyses are presented in Figure 1A and B. In these plots, anole microbiota profiles displayed a gradient that recapitulated the longitudinal transect from Mayagüez (urban) to Maricao (rural), and the microbiota of urban Mayagüez anoles were closer in compositional space to the microbiota of humans from urban areas than were the microbiota of rural anoles. Similarly, the gut microbiota of urban coyotes (Edmonton) were closer in compositional space to those of humans than were those of coyotes from more rural areas (Leduc). Using pairwise beta diversities, we then tested, for both anoles and coyotes, whether the mean beta diversity between urban wildlife and humans was significantly lower than that between rural wildlife and humans (Supplemental Materials). Results indicated that, for both wildlife species, binary Sorensen-Dice microbiota dissimilarity to humans from the United States was significantly reduced in urban settings relative to rural settings (non-parametric *p*-values = 0.001 for each comparison) (Figure 1C, D). Similar results were observed for analyses based on Bray-Curtis dissimilarities (Figure 1—figure supplement 4). In addition, we assessed the degree of compositional convergence of the gut microbiota between wildlife in urban and rural settings with the gut microbiota of humans from rural populations in Malawi and Venezuela. These analyses also supported the conclusion that urban wildlife microbiota have converged compositionally with human microbiota (Figure 1—figure supplement 5). However, wildlife microbiota in urban settings were more compositionally similar to those of urban human populations (USA) (Figure 1C, D) than they were to those of rural human populations (Malawi and Venezuela) (Figure 1—figure supplement 5) (non-parametric *p*-value = 0.001 for each comparison). These analyses further indicated that the gut microbiota of distantly related vertebrates (anoles, coyotes, and humans) have converged compositionally in urban environments.

**Figure 1.**
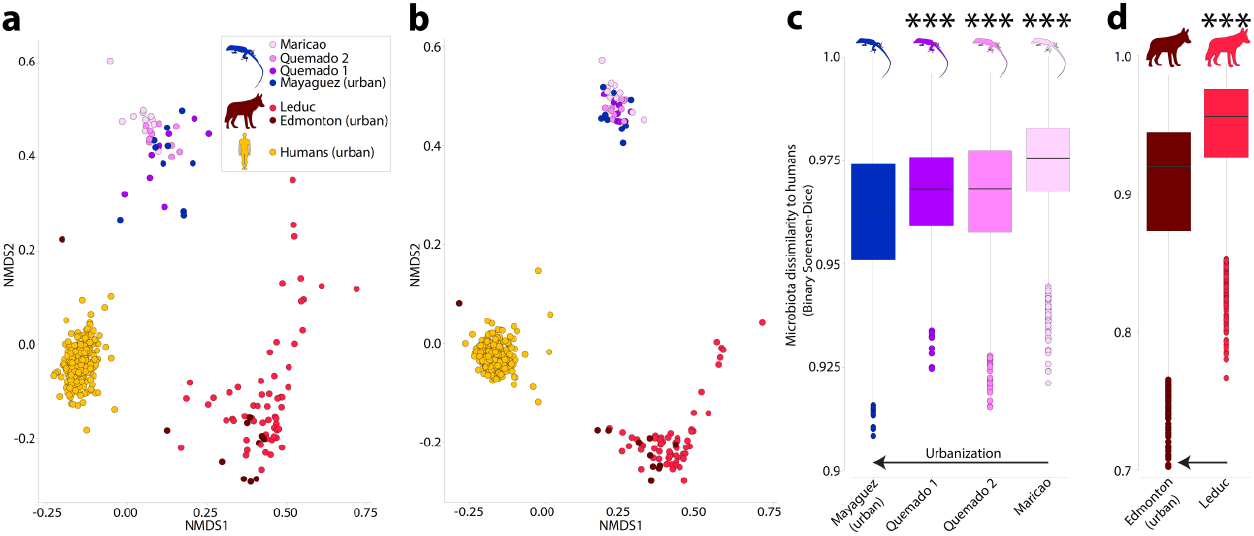
Humanization of urban anole and coyote gut microbiota. Non-metric Multi-dimensional Scaling (NMDS) plots in (a) and (b) show patterns of dissimilarities among anole, coyote, and human (USA adults), gut microbiota profiles. Panels (a) and (b) show NMDS based on Bray-Curtis and binary Sorensen-Dice dissimilarities, respectively. In (a) and (b), each point represents the gut microbiota profile of an individual anole, coyote, or human, as indicated by the key in (a). Boxplots in (c) and (d) show microbiota dissimilarities (binary Sorensen-Dice) between anoles and humans (c) and between coyotes and humans (d). Each box corresponds to comparisons including a single anole or coyote population as indicated. Horizontal arrows indicate gradient of increasing urbanization. Boxplots display median and interquartile range. Significant differences of boxplots compared to urban conspecifics (leftmost boxplot in each panel) based on nonparametric Monte-Carlo permutation tests are denoted with asterisks; *** *p* = 0.001.

To enable identification of the specific ASVs that underlie the observed convergence among human and wildlife gut microbiota in urban settings, we identified all ASVs shared by urban anoles and humans to the exclusion of rural anoles and all ASVs shared by urban coyotes and humans to the exclusion of rural coyotes. A list of these ASVs is presented in Table S4. Three microbial genera— *Clostridium sensu stricto 1, Blautia*, and *Bacteroides*—included >5 ASVs that displayed this pattern. The phylogenetic distributions of these ASVs visualized with Empress (*19*) are presented in Figure 2—figure supplement 1.

To test for differentially abundant ASVs between urban and rural populations, we employed ANCOM (*20*) to compare urban and rural populations of anoles, coyotes, and humans. Results revealed several ASVs that were differentially abundant between urban and rural wildlife populations (Figure 2A and B; Figure 2—figure supplement 2) and between urban and rural human populations (Figure 2—figure supplement 3). We then asked whether the ASVs that were overrepresented in wildlife populations in urban settings relative to conspecifics in rural areas also displayed the same direction of difference in relative abundance between urban and rural human populations. The relative abundance of all ASVs (*Bacteroides vulagtus, Phasocolarctobacterium faecium*, and *Erysipelatoclostridium ramosum*) that were detected in humans and significantly overrepresented in anoles sampled in Mayagüez (urban) relative to anoles sampled in Maricao (rural) are presented in Figure 2C–E. The relative abundances of the ASV (*Ruminococcus gnavus*) that was detected in humans and significantly overrepresented in urban coyotes relative to ruran coyotes is presented in Figure 2F. In every case, the difference in relative abundance between urban and rural wildlife paralleled the difference observed between urban and rural human populations. These ASVs included a predominant human commensal belonging to the genus *Bacteroides* that constituted up to 15% of the gut microbiota in humans from urban environments (Figure 2C). This ASV was also the most significantly overrepresented in urban humans of all ASVs examined (Figure 2— figure supplement 3). ASVs showing parallel shifts in abundance in urban humans and urban wildlife also reached moderately high relative abundances in urban wildlife fecal samples (e.g., ~1%), indicating that these ASVs are unlikely to represent transient members of the gut microbiota. The genera *Bacteroides* and *Ruminococcus*, to which some of the ASVs overrepresented in urban wildlife belong, have been previously established as an indicator taxa of urbanization in humans (*17*, *21*). Furthermore, comparative studies of microbiomes across Earth’s environments have indicated that the human gut is a primary reservoir of these ASVs (*22*). Therefore, humans represent a likely source of these gut microbiota constituents in urban wildlife.

**Figure 2.**
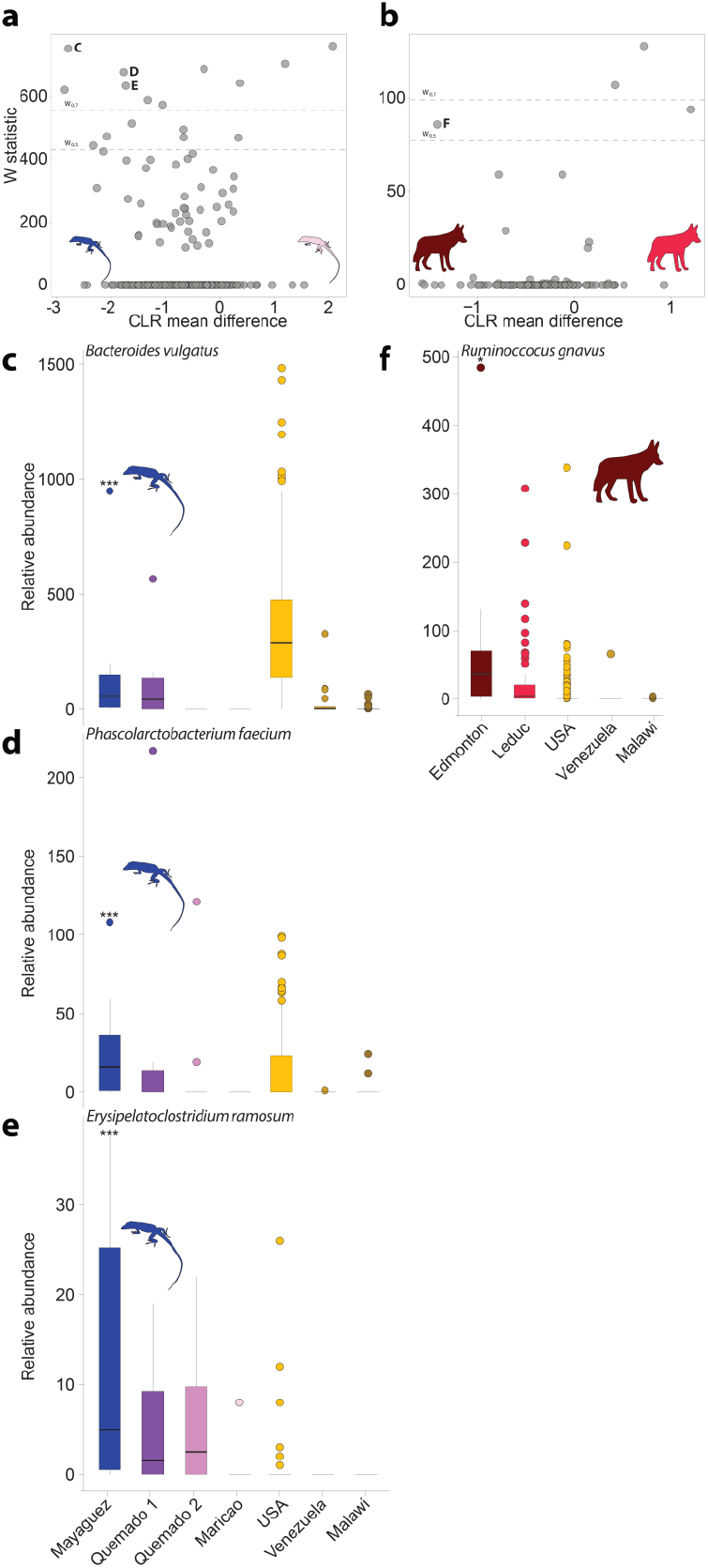
Differentially abundant ASVs between populations in urban and natural settings. Volcano plots display the results of ANCOM tests for ASVs that were differentially abundant between urban and rural populations of anoles (a) and coyotes (b). Each point represents an ASV, with the x-axis denoting the centered log ratio mean difference of the relative abundance of the ASV between urban and rural populations. The y-axis indicates the ANCOM test statistic (W), and horizontal dashed line indicates the significance level W=0.5 and W=0.7. Panel (a) displays results of ANCOM tests comparing Mayagüez and Maricao anoles. Results of ANCOM tests comparing Mayagüez anoles with anoles from the other two sampling locations are presented in Figure 2—figure supplement 2. Boxplots in (c)-(e) show the relative abundances of each of the ASVs that were significantly overrepresented in Mayagüez (urban) anoles relative to Maricao anoles based on ANCOM analyses shown in (a). Boxplots in (f) show the relative abundances of the single ASV that was significantly overrepresented in Edmonton (urban) coyotes relative to Leduc coyotes based on ANCOM analyses shown in (b). Boxplots display median and interquartile range. In (c)-(e), the relative abundances of the ASVs in humans from USA, Venezuela, and Malawi are shown in the rightmost three boxplots in each panel. Note that each ASV displays parallel shifts in relative abundance in urban populations of anoles, coyotes, and humans relative to rural populations of the same host species. Significant differences between ASV relative abundance in wildlife from urban (Mayagüez or Edmonton) and rural (Maricao or Leduc) environments are indicated by asterisks above the boxplot for each urban population; * W > 0.5; *** W > 0.7.

The observation that a set of ASVs found in humans increased in prevalence in urban wildlife is consistent with bacterial spillover from humans into anoles and coyotes in cities. Testing the rates at which bacterial lineages transmit between humans and wildlife in cities will require further strain-level profiling of the gut microbiota of these hosts combined with phylogenetic analyses of strain relationships. A non-mutually exclusive explanation is that parallel dietary shifts select for common sets of ASVs in humans and wildlife. For example, previous work in humans has shown that the relative abundance of *Bacteroides* is associated with diets high in animal fat and protein (*23*). It is possible that dietary differences between urban and rural wildlife populations may contribute to the observed gut microbiota convergence with humans in cities. Our results motivate future profiling of diets (e.g., through metabarcoding) and gut microbiota composition to test whether shared dietary differences between urban and rural populations drive convergence of the gut microbiota.

Overall, this study demonstrates the convergence of gut microbiota composition among distantly related vertebrates living in urban environments. Interestingly, this convergence was evident despite the fact that the human and wildlife populations that we examined resided in different cities throughout North America, indicating parallel effects of urbanization on humans and wildlife independent of geographic location. Close contact among hosts of the same species can generate social microbiomes—microbial metacommunities formed by conduits of microbial transmission along host social networks (*24*). Previous studies have also indicated that transmission of the gut microbiota can occur between species of mammalian wildlife when they come into close contact, such as predator-prey interactions (*2*). Our results suggest that urban environments host gut microbiota metacommunities of vertebrate species as distantly related as reptiles and mammals, implying the existence of gut microbial transmission routes among humans and diverse wildlife in cities.

## Supplemental Methods

### Sample collection

Fecal samples were collected from anoles (*Anolis cristatellus*) in four sampling locations shown in Figure 1A. Animals were caught with lassos and placed into clean plastic bags for fecal collection. All fecal material was snap frozen upon collection and stored at −80C. Sample sizes were determined by catch success over a nine-day period from July 14, 2019 to July 23, 2019.

### DNA Extractions, library preparation and sequencing

DNA was extracted from all anole fecal pellets with a bead beating procedure based on the Qiagen PowerLyzer kit. The V4–V5 region (515F 926R primer pair) of 16S rRNA gene was amplified from all DNAs in duplicate with the high fidelity Phusion polymerase as described by Comeau et al., 2017. Following library preparation, 16S rRNA gene libraries were pooled in equimolar amounts and sequenced on a single lane of Illumina MiSeq using 300 + 300 bp paired-end V3 chemistry following protocols of (*25*).

### Quality filtering, sequence processing, and taxonomic assignments

Raw fastq files generated from anole fecal microbiota libraries were uploaded to the qiita webserver (qiita.ucsd.edu) and combined with publicly available fastq files containing reads corresponding to 16S rRNA gene V4 sequences from the gut microbiota of adult humans (*17*) and coyotes (*14*). Raw reads were filtered for quality using split libraries and trimmed to a common length of 100bp to enable comparisons across datasets. Amplicon sequence variants were called using deblur as implemented in qiita using default parameters. All ASVs were assigned to taxonomic ranks against the Silva 138 database using the taxonomy command in QIIME2. Samples whose read depths were more than three standard deviations below the mean of the samples’ group or for which >75% of the reads belonged to a single ASV were removed prior to downstream analyses. For each analysis, samples were rarefied to a common depth of 90% of the minimum read depth of samples included in the analysis in order to enable direct comparisons of alpha and beta diversity among samples.

### Beta and alpha diversity analyses

Beta diversities (Binary Sorensen-Dice and Bray-Curtis dissimilarities) were calculated between all pairs of samples and for each individual sample with thein R Phyloseq package (*26*). Alpha diversities (chao1 and shannon) were calculated for each sample and plotted by sampling location in QIIME2. Comparisons of beta diversities between sample groups were conducted with anosim using Kruskal-Wallis tests and PEMANOVA/PERMDISP using the R vegan package. Significance of comparisons of alpha diversities between sample groups was assessed with Kruskal-Wallis tests in QIIME2. Beta dispersion within groups was measured with the R vegan package. Beta diversities were visualized using non-metric multidimensional scaling as implemented in the R phyloseq package.

### Statistical analyses of beta diversity

The hypothesis that the gut microbiota of urban wildlife displayed increased compositional similarity to human gut microbiota relative to the gut microbiota of non-urban wildlife was assessed based on comparisons of Binary Sorensen-Dice and Bray-Curtis dissimilarities with Monte Carlo permutation tests. Specifically, these tests assessed whether beta diversity between urban wildlife (either anoles from Mayaguez or coyotes from Edmonton) and humans was significantly lower than that between non-urban wildlife (anoles from Quemado 1, Quemado 2, or Maricao or coyotes from Leduc) and humans. These tests included 999 permutations of the beta diversity matrices as implemented in make_distance_boxplots.py in QIIME1.

### Phylogenetic analyses of ASVs

A phylogeny of all ASVs was generated using SEPP insertion against the Greengenes 13.8 reference. The ASVs that were shared by urban wildlife and humans but not by non-urban wildlife were identified in R, and the phylogenetic distributions of these ASVs were plotted with empress (*19*).

### Differential abundance testing and visualization

Analysis of compositions of microbiomes with bias correction (ANCOM) (*20*) was employed to test for differentially abundant ASVs between sample groups. For these analyses, sample groups considered were urban versus non-urban anoles, urban versus non-urban coyotes, humans from USA versus humans from Malawi, and humans from USA versus humans from Venezuela. Differentially abundant ASVs were visualized using ggplot. Scripts that were modified for these analyses are available at https://github.com/FrederickHuangLin/ANCOM. The specific scripts used in our analyses are available at https://github.com/briandill2/MicrobiotaUrbanization. Boxplots of ASV relative abundances were made with ggplot. For boxplots of ASV relative abundances, we identified each ASV that was significantly overrepresented in Mayaguez anoles compared to Maricao anoles or significantly overrepresented in Edmonton coyotes compared to Leduc coyotes.

## Supplemental Results

### Statistically significant differences in beta diversity among sample groups

PERMANOVA and PERMDISP analyses indicated significant differences in beta diversity centroids among sample groups. Sample groups included Mayaguez anoles, Quemado 1 anoles, Quemado 2 anoles, Maricao anoles, urban coyotes, peri-urban coyotes, USA humans, Venezuela humans, and Malawi humans. Tests of differences in mean beta diversity between pairs of comparisons of sample groups were conducted with nonparametric tests based on 999 Monte Carlo permutations of the beta diversity matrix as implemented in QIIME. In each pairwise comparison, posthoc Kruskal-Wallis tests of PERMANOVA indicated significant differences among groups (*p* < 0.01 for each comparison). In contrast, no significant differences were observed among sample dispersion based on PERMDISP in 44 out of 44 pairwise comparisons of Sorensen-Dice dissimilarities (*p* > 0.05 for each comparison) and 42 out of 44 pairwise comparisons of Bray-Curtis dissimilarities. The only sample groups displaying significantly different dispersions based on PERMDISP analyses of Bray-Curtis dissimilarities were USA human and Periurban coyotes (*p*-value =1.7e-5) and USA humans and Mayaguez anoles (*p*-value = 0.048).

## Supplemental Figure Legends

**Figure 1—figure supplement 1.**
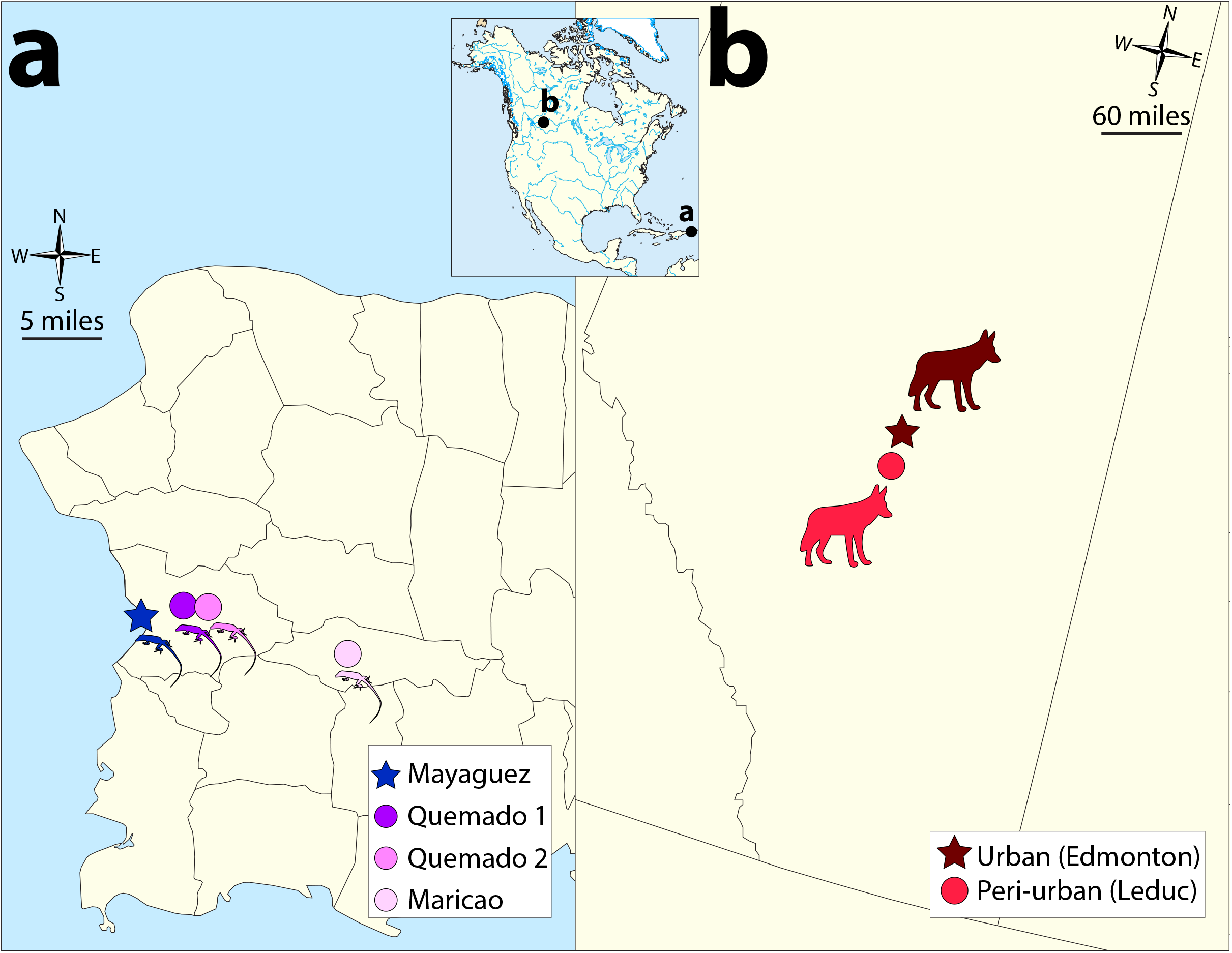
Map of wildlife sampling locations. (a) Circles and star indicate **s**ampling sites of anole populations as indicated by the key. (b) Circle and star indicate sampling sites of coyote populations as indicated by the key. In (a) and (b), stars indicate urban sampling locations and circles indicate rural sampling locations.

**Figure 1—figure supplement 2.**
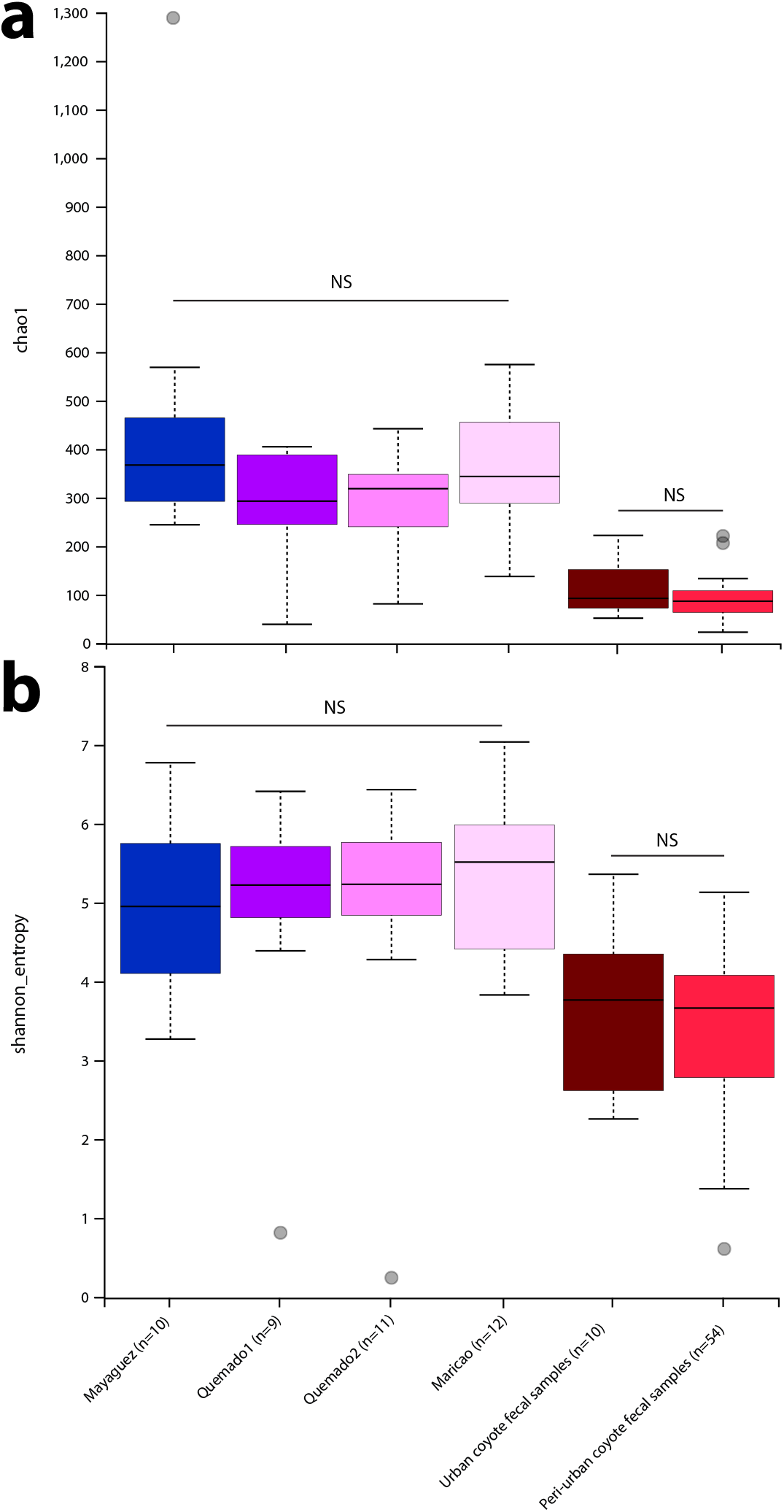
Alpha diversity in gut microbiota of urban and non-urban anoles and coyotes. Boxplots show median and interquartile ranges of alpha diversity estimates for anoles and coyotes. Chao1 and Shannon estimates are shown in (a) and (b), respectively. Significant differences based on Kruskal-Wallis tests are shown; NS *p* > 0.05.

**Figure 1—figure supplement 3.**
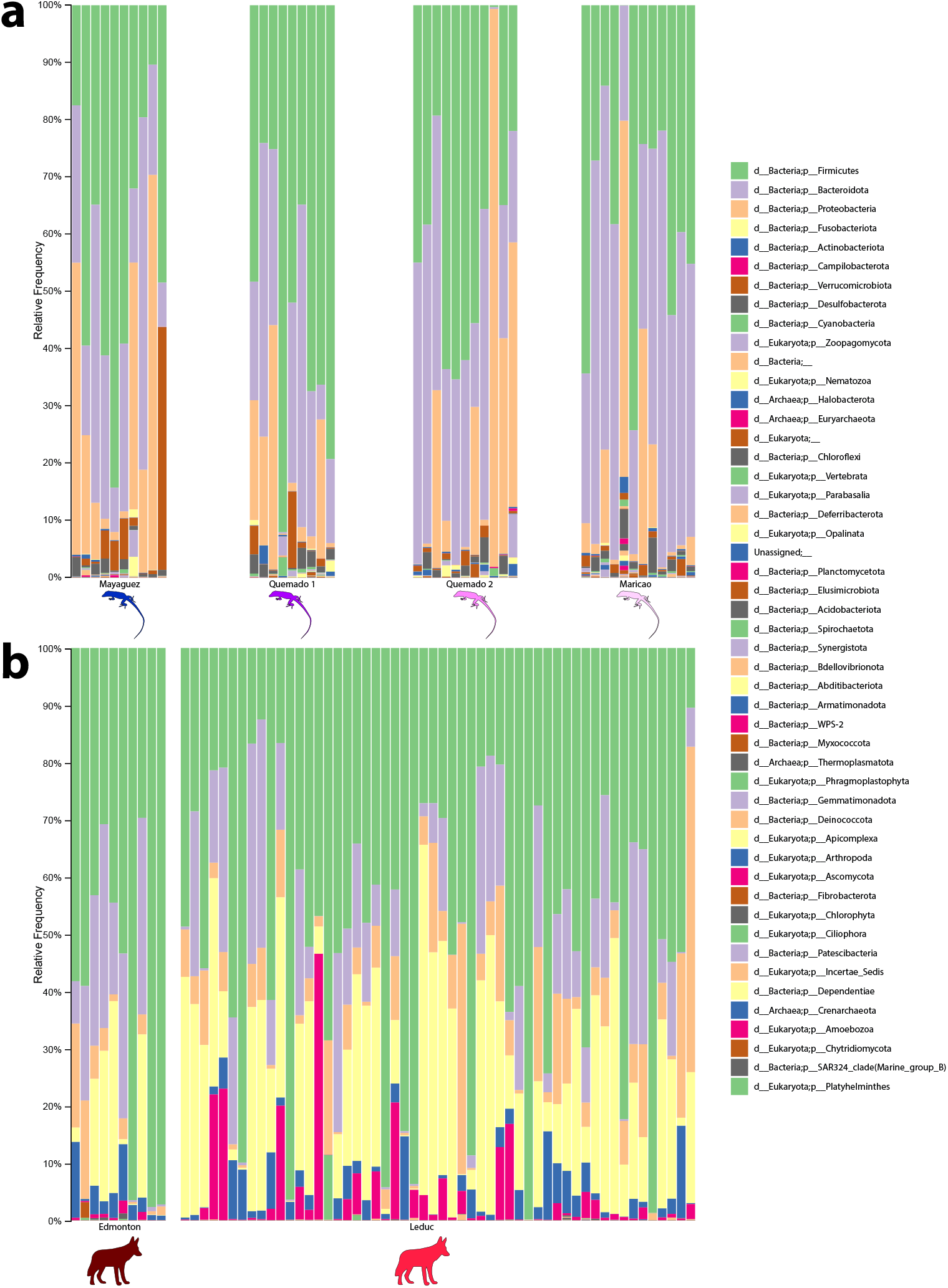
Taxonomic profiles of anole and coyote gut microbiota in urban and non-urban locations. Stacked bar plots display the relative abundances of bacterial phyla in anoles and coyotes from locations shown in Figure 1. Each bar corresponds to a bacterial phylum as indicated by the key.

**Figure 1—figure supplement 4.**
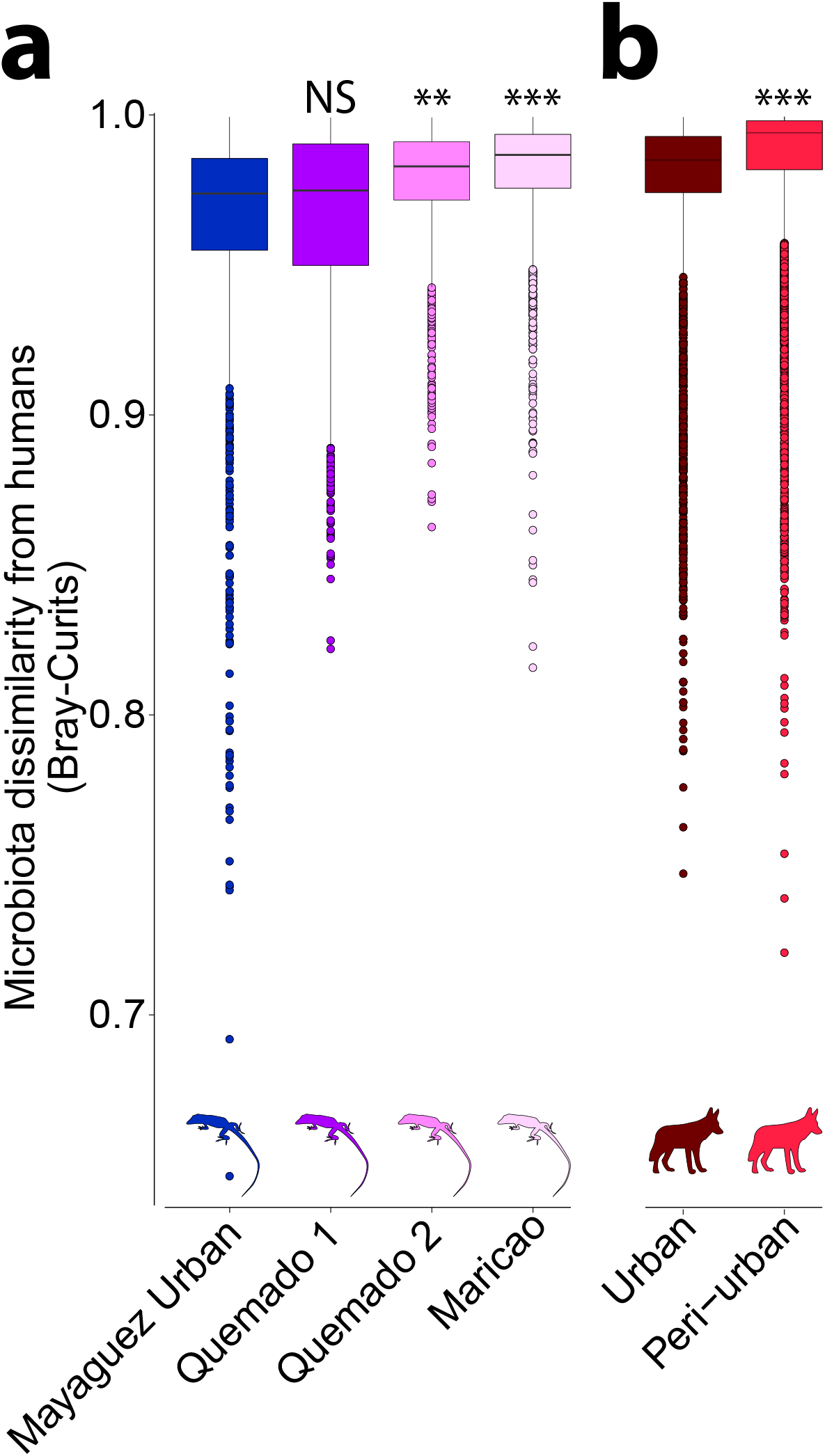
Humanization of urban anole and coyote gut microbiota based on ASV relative abundance profiles. Boxplots show microbiota dissimilarities (Bray-Curtis) between anoles and humans (a) and between coyotes and humans (b). Each box corresponds to comparisons including a single anole or coyote population as indicated, and colors correspond to those in Figure 1. Boxplots display median and interquartile range. Significant differences of boxplots compared to urban conspecifics (leftmost boxplot in each panel) based on nonparametric Monte-Carlo permutation tests are denoted with asterisks; ** *p* < 0.01, *** *p* = 0.001.

**Figure 1—figure supplement 5.**
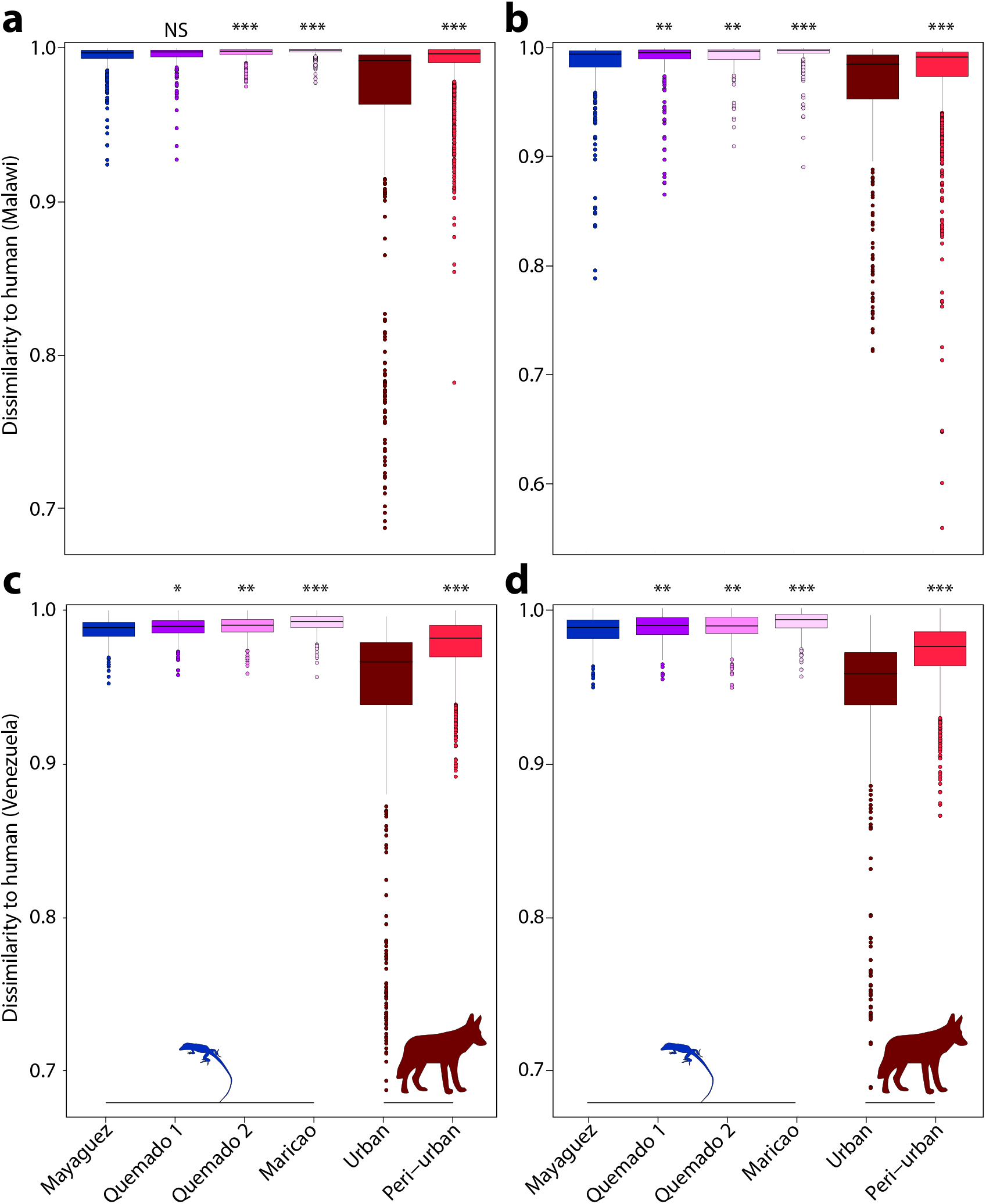
Humanization of urban anole and coyote gut microbiota based on comparisons with humans from Venezuela and Malawi. Boxplots in (a) and (b) show microbiota dissimilarities between anoles and humans from Malawi based on Bray Curtis (a) and binary Sorensen-Dice (b) dissimilarities. Boxplots in (c) and (d) show microbiota dissimilarities between anoles and humans from Venezuela based on Bray Curtis (c) and binary Sorensen-Dice (d) dissimilarities. Each box corresponds to comparisons including a single anole or coyote population as indicated, and colors correspond to those in Figure 1. Boxplots display median and interquartile range. Significant differences of boxplots compared to urban conspecifics (leftmost boxplot in each panel) based on nonparametric Monte-Carlo permutation tests are denoted with asterisks; NS *p* > 0.05, * *p* < 0.05, ** *p* < 0.01, *** *p* = 0.001.

**Figure 2—figure supplement 1.**
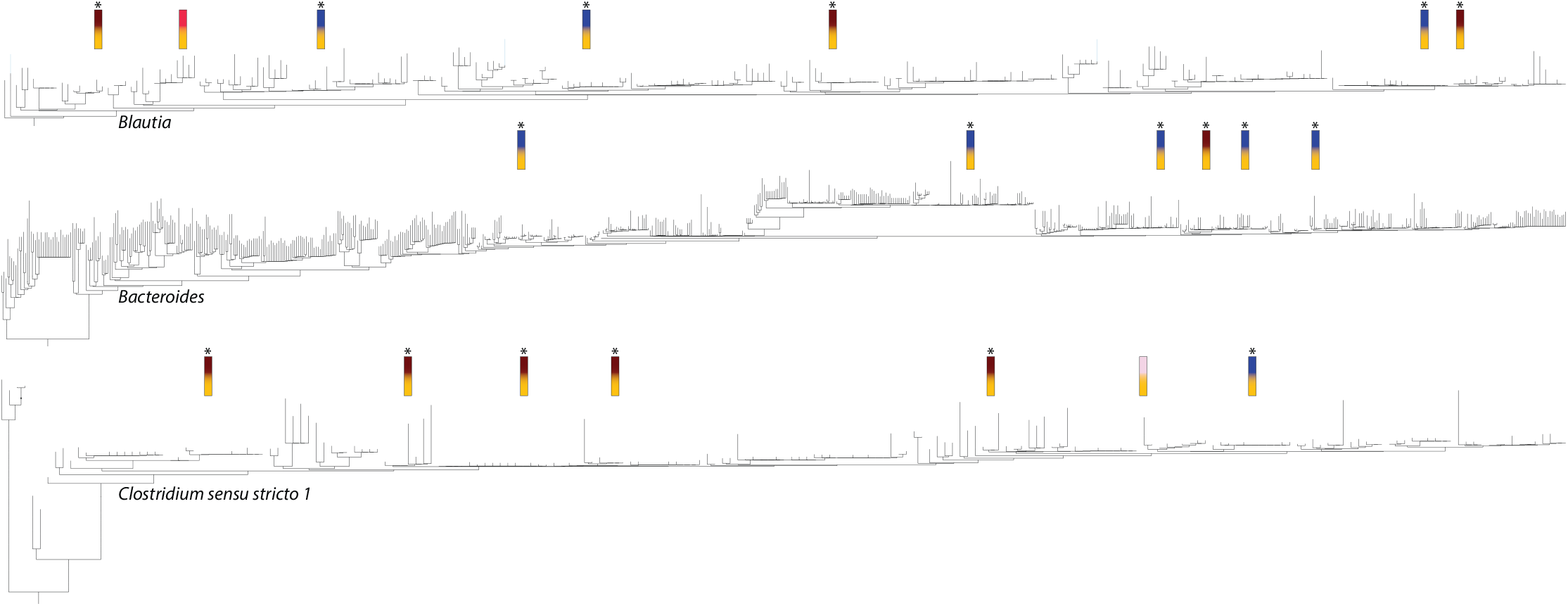
Phylogenetic distribution of ASVs shared by humans and urban wildlife. Phylogenies show relationships among ASVs within microbial genera (*Blautia, Bacteroides*, and *Clostridium sensu scrito 1*) for which at least 5 ASVs were shared exclusively by humans and urban wildlife (to the exclusion of rural wildlife). Vertical bars mark the ASVs that were either shared by urban wildlife and humans to the exclusion of rural wildlife or shared by rural wildlife and humans to the exclusion of urban wildlife. Colors of vertical bars indicate the host populations in which the ASVs were detected as indicated in the key in Figure 1A. Vertical bars with asterisks correspond to the 18 ASVs that displayed a distribution across host populations indicative of humanization, whereas vertical bars without asterisks correspond to the two ASVs that displayed the opposite distribution (i.e., shared by rural wildlife and humans to the exclusion of urban wildlife).

**Figure 2—figure supplement 2.**
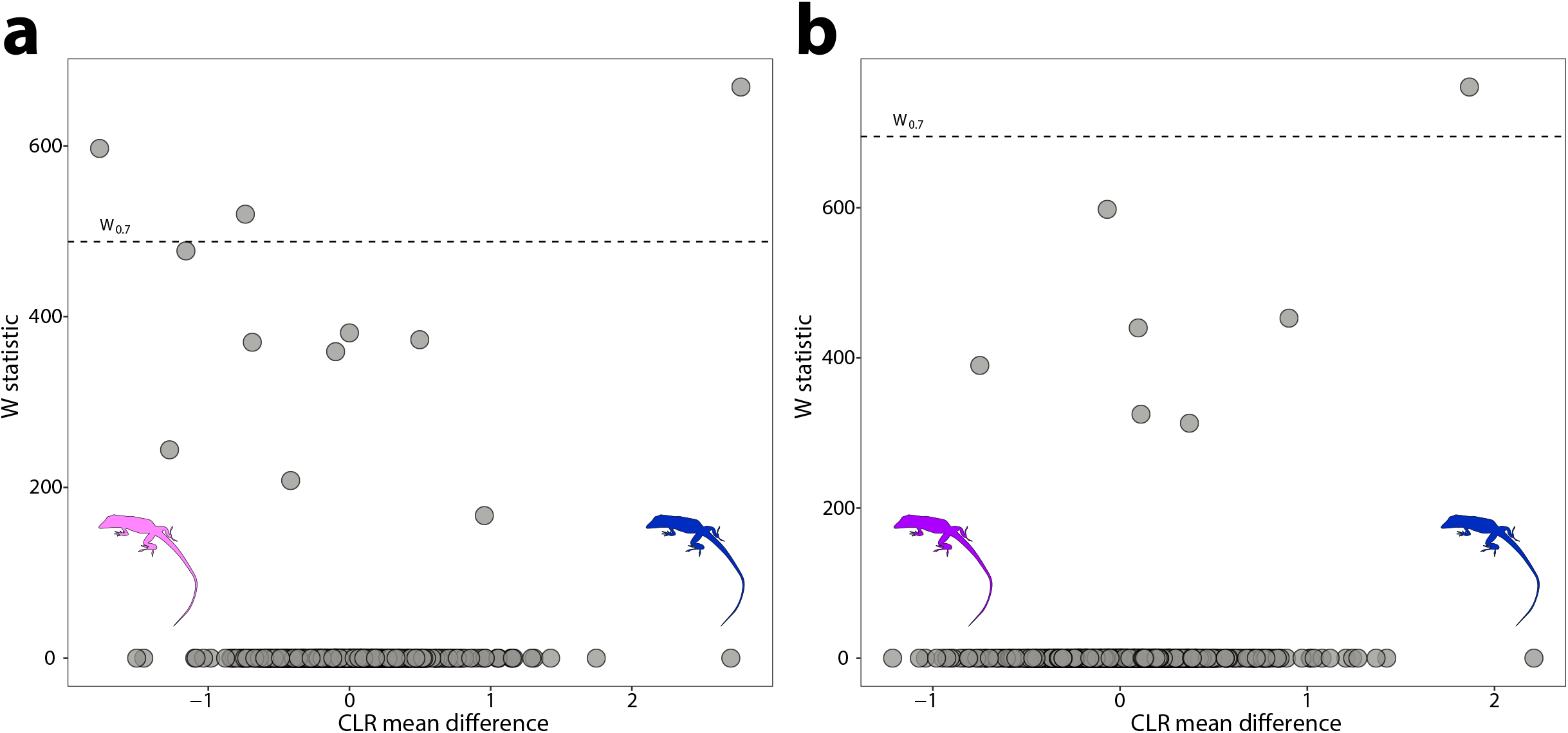
Differentially abundant ASVs between anole populations in urban and natural settings. Volcano plots display the results of ANCOM tests for ASVs that were differentially abundant between urban and natural populations of anoles from (a) Mayaguez and Quemado 1 and (b) Mayaguez and Quemado 2. Each point represents an ASV, with the x-axis denoting the centered log ratio mean difference of the relative abundance of the ASV between urban and natural populations. The y-axis indicates the ANCOM test statistic (W), and horizontal dashed lines indicate the significance levels W=0.5 and W=0.7.

**Figure 2—figure supplement 3.**
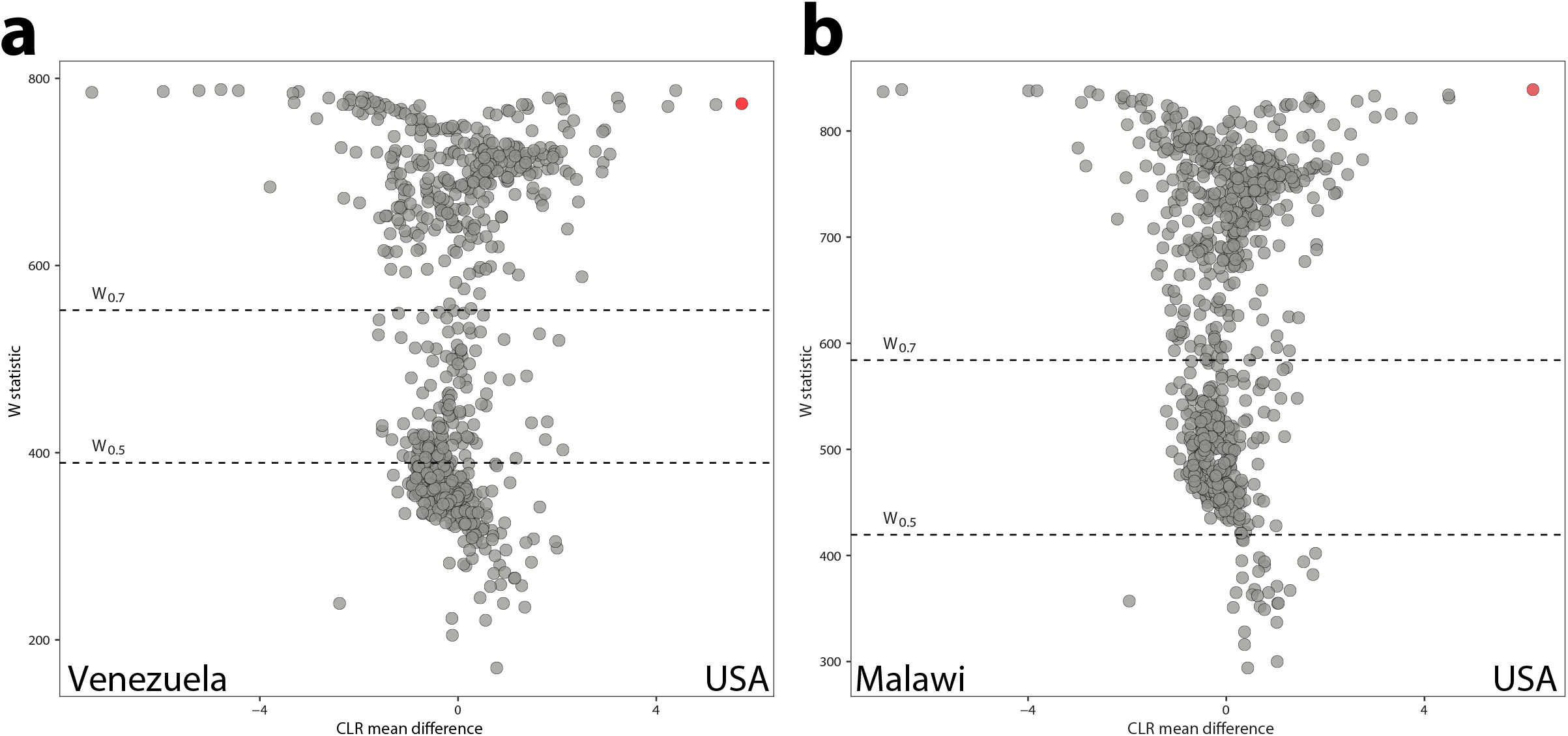
Differentially abundant ASVs between human populations. Volcano plots display the results of ANCOM tests for ASVs that were differentially abundant between urban (USA) and non-urban (Venezuela and Malawi) populations of humans. Comparisons between USA and Venezuela populations are show in (a), and between USA and Malawi populations in (b). Each point represents an ASV, with the x-axis denoting the centered log ratio mean difference of the relative abundance of the ASV between urban and natural populations. Red points indicate the *Bacteroides* ASV presented in Figure 2. The y-axis indicates the ANCOM test statistic (W), and horizontal dashed lines indicate the significance levels W=0.5 and W=0.7.

## Supplemental Tables

**Table S1. Metadata for all samples.**

**Table S2. ASV relative abundances across all samples.**

**Table S3. Taxonomic assignments of all ASVs.**

**Table S4. ASVs shared by urban wildlife and humans but not by rural conspecific wildlife.**

